# Ranking Reprogramming Factors for Directed Differentiation

**DOI:** 10.1101/2021.05.14.444080

**Authors:** Jennifer Hammelman, Tulsi Patel, Michael Closser, Hynek Wichterle, David Gifford

## Abstract

Transcription factor over-expression is a proven method for reprogramming cells to a desired cell type for regenerative medicine and therapeutic discovery. However, a general method for the identification of reprogramming factors to create an arbitrary cell type is an open problem. We examine the success rate of methods and data for directed differentiation by testing the ability of nine computational methods (CellNet, GarNet, EBSeq, AME, DREME, HOMER, KMAC, diffTF, and DeepAccess) to correctly discover and rank candidate factors for eight target cell types with known reprogramming solutions. We compare methods that utilize gene expression, biological networks, and chromatin accessibility data to identify eight sets of known reprogramming factors and comprehensively test parameter and pre-processing of input data to optimize performance of these methods. We find the best factor identification methods can identify an average of 50-60% of reprogramming factors within the top 10 candidates, and methods that use chromatin accessibility perform the best. Among the chromatin accessibility methods, complex methods DeepAccess and diffTF are more likely to consistently correctly rank the significance of transcription factor candidates within reprogramming protocols for differentiation. We provide evidence that AME and DeepAccess are optimal methods for transcription factor recovery and ranking which will allow for systematic prioritization of transcription factor candidates to aid in the design of novel reprogramming protocols.

## Introduction

Our ability to precisely discover, define, and characterize cell types has improved with the advent of new molecular technologies in the past decade (Pellegrino et al. 2016; Habib et al. 2016; Corces et al. 2017; Rai et al. 2020). Advances in single cell sequencing methods has even made it possible to define individual cell types by their expression (Sasagawa et al. 2013; Dixit et al. 2016; Angermueller et al. 2016; Grün et al. 2015; Pijuan-Sala et al. 2019) and chromatin accessibility profiles (Lake et al. 2018; Pijuan-Sala et al. 2020; Satpathy et al. 2019). The identification and characterization of cell types in health and disease has further illuminated the potential of regenerative medicine that would be enabled by the ability reprogram cells into arbitrary types. In order to facilitate the identification of reprogramming factors that can be deployed any cell type of interest, we have systematically compared computational methods to identify the most robust means for identification of potential reprogramming factors from gene expression and chromatin accessibility data.

Reprogramming strategies typically either employ small-molecules that target signaling pathways or transcription factor-based reprogramming (Wichterle et al. 2002; Marson et al. 2008; Ichida et al. 2009). Transcription factor over-expression has been a successful method for reprogramming cells such as fibroblasts or stem cells into many specialized cell types (Oh and Jang 2019). Transcription factor protocols are also advantageous as they can decrease protocol time and increase efficiency (Mazzoni et al. 2013). Identification of reprogramming transcription factors has generally combined expert knowledge and large-scale screens to test many possibilities and experimentally identify a set of key regulatory factors. For example, the four transcription factors that induce pluripotency were discovered using a candidate set of 24 transcription factors curated from the literature that were narrowed down to four factors using successive leave-one-out experiments to reprogram fibroblasts into stem cells (Takahashi and Yamanaka 2006).

The ability to easily generate gene expression and chromatin accessibility data from primary cells has led to the development of several computational methods that can potentially discover reprogramming factors even in the absence of extensive developmental studies. However, the performance of these methods has not been systematically compared using the same source data. Here we comprehensively and uniformly evaluate nine reprogramming factor discovery methods: CellNet (Cahan et al. 2014; Radley et al. 2017; Morris et al. 2014), GarNet (Tuncbag et al. 2016; Kedaigle and Fraenkel 2018), EBSeq (Leng et al. 2013), AME (Whitington et al. 2011), DREME (Bailey 2011), HOMER (Heinz et al. 2010), KMAC (Guo et al. 2018), DeepAccess (Hammelman et al. 2020), and diffTF (Berest et al. 2019) on their ability to recover and rank eight known sets reprogramming transcription factors.

Gene expression data have been a long-standing basis for the identification of reprogramming factors (Heinäniemi et al. 2013; Roost et al. 2015; Lang et al. 2014; D’Alessio et al. 2015; Rackham et al. 2016). The differential expression of transcription factors in a target cell type is an indicator of their potential as reprogramming factors. However, there may be both biological and experimental confounders in using expression data for identification of reprogramming factors. For example, it has been shown that sensory neurons of the dorsal root ganglia co-express many transcription factors as they are being specified but change their expression patterns to only express select transcription factors as they mature (Sharma et al. 2020). Additionally, gene expression data does not provide information about whether proteins are present and actively binding DNA to control transcription. When gene expression datasets generated in different studies are used together, experimental confounders such as the differences in measurements that result from nuclear or whole cell mRNA, or using different RNA amplification methods for sequencing. Network methods such as CellNet (Cahan et al. 2014; Morris et al. 2014; Radley et al. 2017) and Inferelator (Bonneau et al. 2006; Miraldi et al. 2019) have been shown to better prioritize transcription factor candidates through the incorporation of gene expression and transcription factor-gene interaction networks, but rely on massive repositories of RNA-seq data measured from perturbed cells to confidently learn these biological networks. As a result, these network-based methods are not generally applicable to novel data from a small number of experiments.

Chromatin accessibility-based methods for identifying candidate reprogramming factors, such as HOMER, AME, DREME, and KMAC characterize over-represented transcription factor binding motifs in accessible chromatin in a target cell type or learn to predict the relationship between chromatin accessibility and DNA sequence (DeepAccess) or measure the differential accessibility of transcription factor sites (diffTF). ATAC-seq can measure the accessibility of chromatin in small cell populations and thus can be used with cells derived from *in vivo* samples, while DNase-seq requires a larger number of cells (Buenrostro et al. 2015; Liu et al. 2019). DNA sequences that are over-represented in accessible genomic regions typically contain informative transcription factor binding motifs. Known motif enrichment or *de novo* motif discovery methods are commonly applied to ATAC-seq data, and require the selection of differentially accessible regions in the starting and target cell types. In using these methods, parameters such as choice of accessibility or histone mark genomic data, number and choice of genomic regions from target cells, and choice of background sequences using shuffled or natural genomic sequences must be chosen carefully to the generate the best quality results. GarNet (Tuncbag et al. 2016; Kedaigle and Fraenkel 2018) combines ATAC-seq and RNA-seq with the goal of identifying transcription factors which are likely to control differential gene expression, though methods that combine gene expression and chromatin accessibility may be subject to the same biological and experimental confounders faced by methods which rank transcription factors using differential gene expression.

Overall, we find that methods that use chromatin accessibility have superior reprogramming factor discovery performance when compared with gene expression methods. We identify optimal accessible region selection strategies for sequence-based methods and using these optimal strategies, we find that AME has the most robust performance for transcription factor recovery, but DeepAccess is the best performer when ranking the significance transcription factors within reprogramming protocols. We also find that histone mark and EP300 annotation do not significantly improve transcription factor recovery which suggests accessibility alone is sufficient to identify reprogramming factors for new target cell types.

## Results

### Chosen factor discovery methods use gene expression or chromatin accessibility

The nine reprogramming factor discovery methods we evaluated used gene expression (EBSeq, CellNet), chromatin accessibility (DREME, AME, Homer, KMAC, diffTF, DeepAccess), or a combination of the two (GarNet) to identify transcription factors as reprogramming candidates based on their popularity and spanning a diverse set of methodological approaches for reprograming factor discovery. We evaluated the methods on their ability to identify transcription factors that reprogram cells from a starting cell type (stem cells or fibroblasts), to eight possible target cell types (induced pluripotent stem cells, skeletal muscle cells, cardiomyocytes, definitive endoderm cells, hepatocyte cells, pancreatic beta cells, dopaminergic midbrain neurons, or spinal motor neurons) (Figure 1A). We started with RNA-seq and ATAC-seq data from primary cells with the exception of definitive endoderm where the endoderm cells were differentiated with small molecules (Cernilogar et al. 2019), and the basis of our evaluation was the reproduction of known reprogramming factors for each target cell type (Figure 1B). Both RNA-seq and ATAC-seq was collected from the same lab for each cell type with the exception of (Table S1) stem cells where our ATAC-seq and RNA-seq were from different sources. We uniformly processed the RNA-seq and ATAC-seq data (Methods).

**Figure 1.**
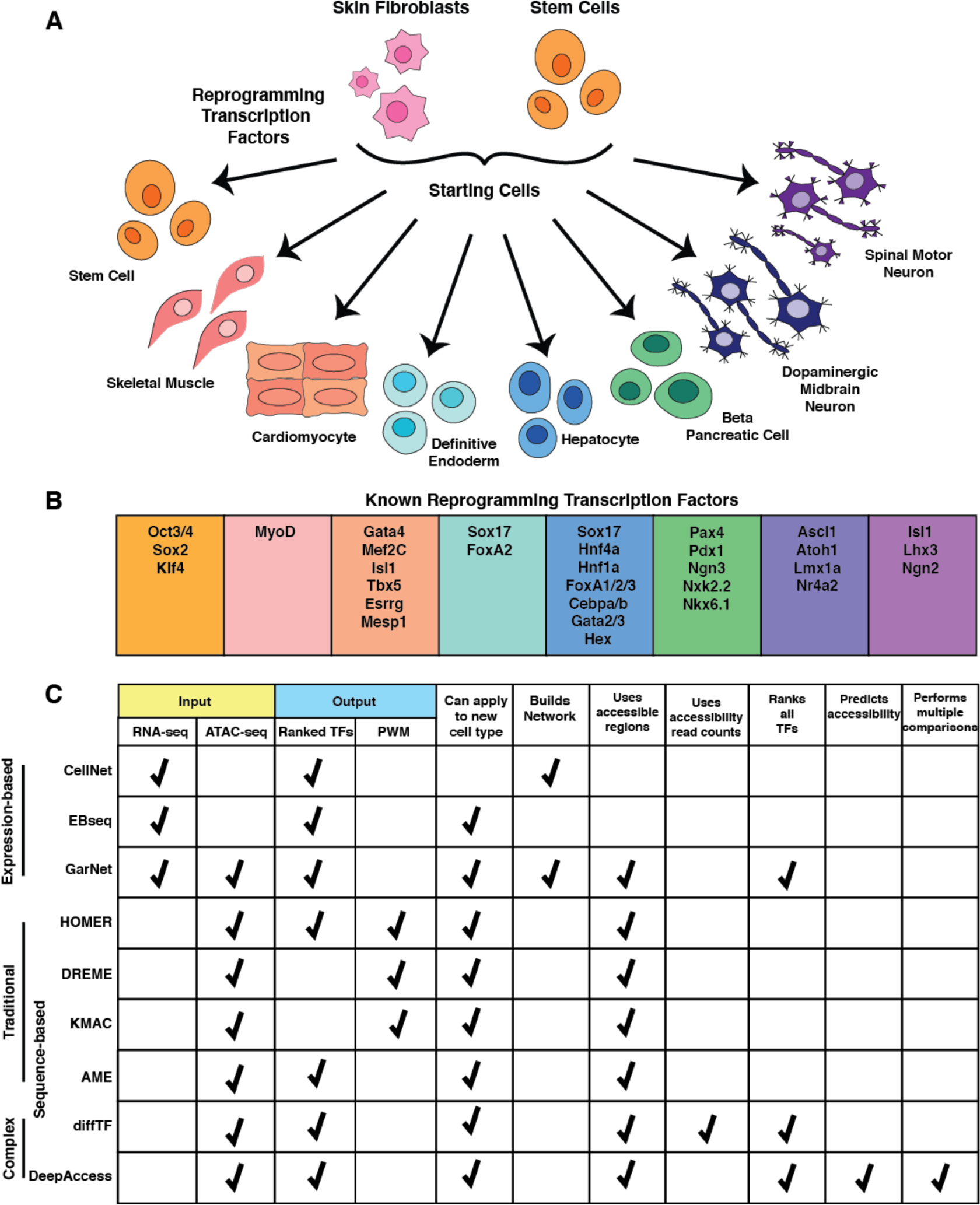
Identifying transcription factors that reprogram starting cells to target cell types. A) Two common starting cell types, fibroblasts (skin cells) and pluripotent stem cells can be reprogrammed to 8 cell types through over-expression of reprogramming transcription factors. B) Transcription factors that have been previously implicated in reprogramming protocols. C) Methods for identifying reprogramming transcription factors from gene expression (RNA-seq) or chromatin accessibility (ATAC-seq). Output of these methods is either a ranked list of transcription factors or a rank representation of DNA binding domain of a transcription factor by a PWM. Most methods can be readily applied to new data, except CellNet which requires many samples from the same cell type to build cell type-specific regulatory networks. All methods that use chromatin accessibility data take in accessible regions, typically generated from a peak calling algorithm such as MACS2 as input. In contrast to more traditional approaches for identifying transcription factors, diffTF and DeepAccess both model more complex regulation by taking into account read counts, and DeepAccess can also predict accessibility of DNA sequences unseen in the genome for other downstream analyses such as cross-species prediction.

We used EBSeq and CellNet as our methods for discovering reprogramming factors from gene expression data. EBSeq (Leng et al. 2013) ranks transcription factor differential expression between a starting and target cell type (Figure 1C). CellNet (Cahan et al. 2014; Morris et al. 2014; Radley et al. 2017) ranks candidate factors based on their importance in a cell type specific regulatory network derived from perturbation-based gene expression datasets. We attempted to train a new CellNet model using our data from the replicated RNA-seq samples from our eight target cell types, but found the experimental data did not represent sufficiently informative perturbations to build cell type specific regulatory networks. Thus, we evaluated CellNet for 5 cell types (stem cell, hepatocyte, cardiomyocyte, skeletal muscle) with preexisting regulatory networks. Consequently, CellNet did not fit our criteria for methods which can be used to predict reprogramming transcription factors from new, perhaps difficult to collect, cell types.

We examined if the combination of differential gene expression and chromatin accessibility could improve reprogramming factor prediction with GarNet (Tuncbag et al. 2016; Kedaigle and Fraenkel 2018). Using a list of transcription factor putative binding sites, which can be derived from motif scanning or ChIP-seq, GarNet assigns binding sites that are present in accessible regions to their closest gene within a constrained distance. It then uses transcription factor scores per genomic site (such as ChIP binding signal or the strength of a motif match) to train a regression model to predict differential gene expression. The weights associated with a particular motif then indicate its potential importance in driving differential gene expression.

Finally, we explored methods for transcription factor motif discovery from chromatin accessibility data. We selected methods that are both widely adopted and varied in approach. From the MEME suite, DREME (Bailey 2011) performs *de novo* motif discovery by first identifying seed sequences as the top 100 significantly enriched words relative to background sequences, then performs a beam search based on the seed sequences to generalize words to PWMs. AME (Machanick and Bailey 2011) performs discriminative motif enrichment from an existing database of motifs represented by PWMs. HOMER (Heinz et al. 2010) first identifies the most globally enriched oligos (similar to DREME) then transforms them into PWMs which get further optimized with a sensitive local optimization algorithm. KMAC (Guo et al. 2018) also performs *de novo* motif discovery, but uses a *k-mer* representation of DNA binding motifs that was shown to better represent actual DNA binding sites by including features such as flanking nucleotides. For the *de novo* motif discovery methods (DREME, HOMER and KMAC), their output is the form of a list of position weight matrices representing enriched DNA binding motifs. To link this to transcription factor activity, we used Tomtom to determine whether each PWM significantly matched one or more known transcription factor motifs (Methods). As motif discovery methods rely on input choices that can affect their performance, we extensively tested (1) the number of accessible regions input, (2) if differential regions were from the starting cell type or the top most accessible regions, and (3) the choice of background sequences.

In addition to traditional motif discovery methods, we applied two complex methods for evaluating motif enrichment, diffTF and DeepAccess. We define complex methods to be those that incorporate read count information or model more complex regulatory grammars. Both complex methods also require access to high-performance computing. diffTF (Berest et al. 2019) uses the accessibility sequencing read counts within putative transcription factor binding sites (either from motif instances or ChIP-seq) to estimate the fold change in accessibility between two conditions for a given transcription factor. We also tested a deep learning estimate of transcription factor activity using DeepAccess, an ensemble of convolutional neural networks trained to predict chromatin accessibility across multiple cell types (Hammelman and Gifford 2021; Hammelman et al. 2020). In previous work, we found that DeepAccess was successful in identifying DNA sequences driving differential accessibility between stem cells and definitive endoderm, as measured by a high-throughput reporter assay for chromatin accessibility (Hammelman et al. 2020). In order to rank transcription factors with DeepAccess, we utilize the Differential Expected Pattern Effect (Hammelman and Gifford 2021) to estimate transcription factor impact on chromatin accessibility using computational DNA sequence perturbation, where we simulate *in silico* an experiment where the DNA binding motif for a given transcription factor is inserted at a previously closed genomic locus. To rank transcription factors as reprogramming candidates, we estimate the Differential Expected Pattern Effect of these transcription factor motifs between the starting and target cell types.

### A consensus database of mouse transcription factor motifs simplified evaluation

We used a custom clustered database of 107 mouse transcription factor motifs to simplify downstream analysis with transcription factor families that share highly similar motifs. Using the HOCOMOCOv11 database (Kulakovskiy et al. 2018) of 356 mouse transcription factor motifs, we computed the pairwise similarity between motifs using Tomtom (Gupta et al. 2007). We then applied Pearson’s correlation and affinity propagation clustering to obtain motif clusters. Affinity propagation clustering is a message passing algorithm that assigns a representative data point to each cluster, which can be weighted to select for ideal characteristics (Frey and Dueck 2007). In this case, the similarity metric was based on the information content of the motif. This resulted in 107 transcription factor motifs representing major families of transcription factors (Figure S1).

### Optimization of accessible region selection for the AME, HOMER, DREME, and KMAC methods

In order to identify transcription factors motifs that are enriched in accessible genomic regions of target cell types, motif discovery algorithms typically compare the prevalence of motifs in sets of positive (target) vs. negative (background) sequences. In order to identify the most effective parameters for motif discovery, we examined three attributes of input regions: 1) for positive sequences, we used either the most significant accessible regions ranked by MACS2 in the target cell type or the most significant accessible regions that are accessible in the target cell type and not accessible in the starting cell type of stem cells or fibroblasts, 2) for negative background sequences we used randomly shuffled positive sequences, enhancers shared across multiple cell types, or GC-content-matched randomly sampled genome sequences, and 3) the number of significant accessible regions to use as input (Figure 2A). For our enhancer background sequences, we used Mouse ENCODE project candidate enhancer annotations from 18 tissues, where enhancers were defined by both the presence of H3K4me1 and absence of H3K4me3 (Shen et al. 2012). To be included in our background sequences, the enhancer had to be present in 15 out of 18 tissues resulting in a total of 2,609 general enhancers. For GC percentage content-matched genome-sampled sequences, we used HOMER to generate sequences then ran AME, DREME, or KMAC using these sequences as background. Over all methods and input attributes, we ran a total of 150 experiments for transcription factor recall over our eight target cell types. Based on the area under the transcription factor recall curve within the top 10 ranked motifs averaged over the eight target cell types, we selected optimal strategies for each motif discovery method (Figure 2B; Table S2-3).

**Figure 2.**
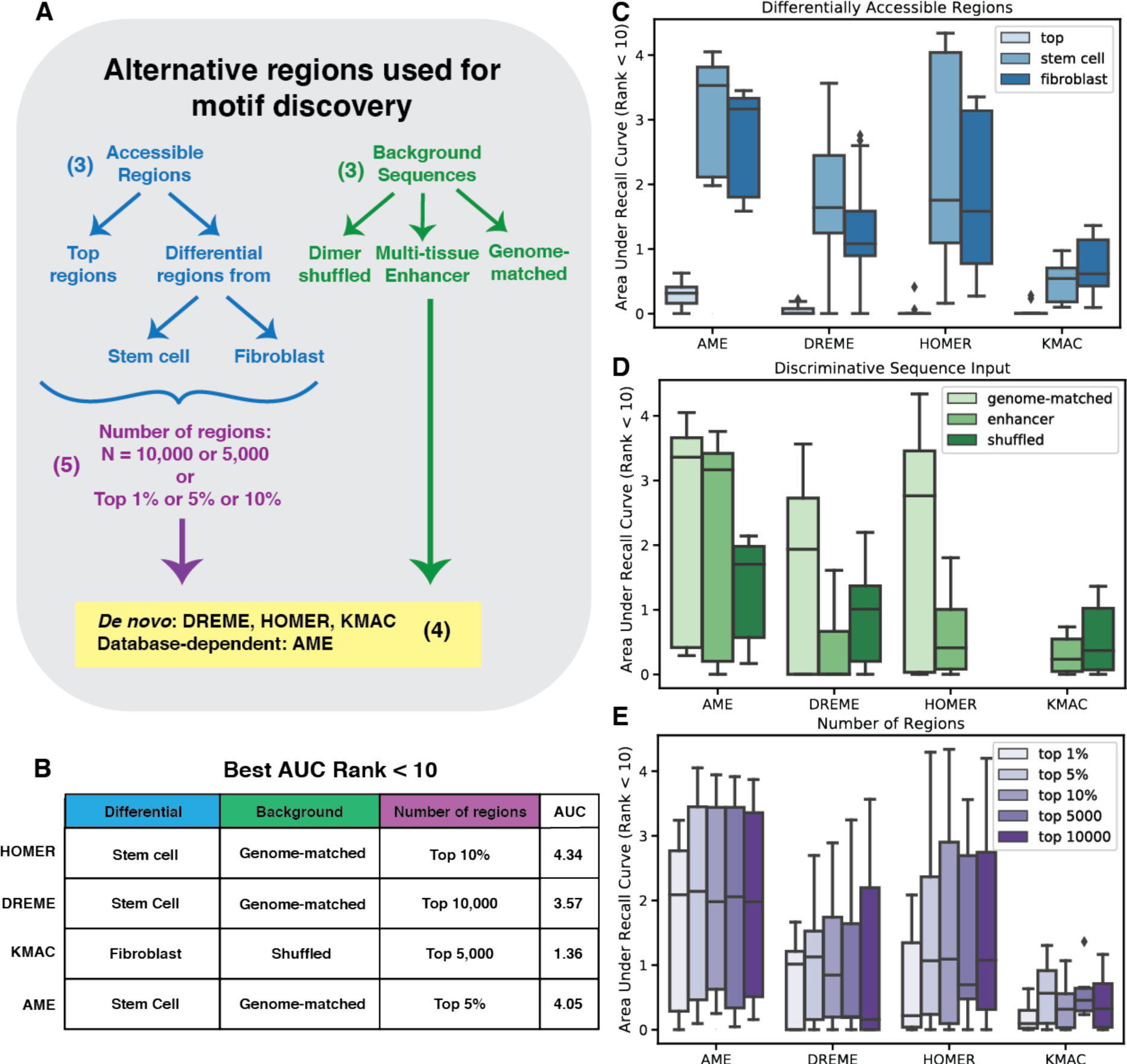
Selection of genomic regions impacts traditional DNA sequence-based methods for identification of transcription factors from chromatin accessibility. A) The 3 axes for selection of genomic regions are choice of top regions or regions that are target cell type-specific relative to starting cell type, choice of background sequences for discriminative motif discovery, and number of regions. For choice of background sequence, we chose to compare universal enhancer sequences to the method default, which for a majority of methods is di-nucleotide shuffled controls, except for HOMER which samples nucleotide-content matching sequences from the genome. B) For each traditional method, the best set of genomic regions are chosen based on transcription factor recovery of all 8 cell types within in the top 10 ranked transcription factor motifs. For all methods, the best performing region selection used differential regions relative to some starting cell type and not top regions. The best performer for each method also used at least 5,000 or 5% of sequences. C) Overall reprogramming factor recovery for all region selections separated by choice of differentially accessible or top regions shows that using differential regions improve transcription factor recovery for all methods. D) Choice of default over enhancer background sequences improves recovery for DREME and HOMER methods but has little impact for AME and KMAC. E) Number of regions has impact on recovery for AME, with more regions leading to more recovery, while other method performance appears to be less impacted by number of regions.

We found that all methods were improved by the use of regions that were accessible in the target cell type and not in stem cells or fibroblasts, with stem cells as a preferred starting cell type for 3 out of 4 methods (Figure 2C; Table S3). We then investigated the performance given the choice of background sequences for discriminative motif discovery. GC%-matched genome-sampled background sequences improved performance for AME, DREME, and HOMER over shuffled or multi-tissue enhancer sequences. Due to the memory required for oligo-based motif discovery, we were unable to consistently run KMAC using the 40,000 – 50,000 genome-matched background sequences generated by HOMER. One notable difference was that enhancer-based background sequences performed well for AME but not for DREME or HOMER. One possible reason is the first seed oligo selection step that is common to both DREME and HOMER that may be affected when the number of background sequences is significantly different from the input sequences. In contrast to other input attributes, the number of input sequences had little effect on performance for most methods, except using the top 1% of sequences appeared to be too few to robustly recover transcription factors (Figure 2E). Overall, it seemed that using 5,000-10,000 input sequences yielded robust success across all methods, as taking the top 5-10% of regions typically also fell in this range (Figure 2B).

### Addition of histone mark and EP300 information does not impact performance of sequence-based methods

One question important for prioritizing experiments for stem cell biologists is whether the addition of histone mark or EP300 data which further implicates genomics regions as cell typespecific cis-regulatory elements would improve performance for transcription factor recovery. Such histone marks have helped to improve enhancer recall in previous work (Fu et al. 2018). Since matched histone mark data was not available for all cell types, we focused on the liver where we could use ATAC-seq, H3K27ac, H3K4me3, EP300, and H3K4me1. The area under the recall curve (Figure 3A) and the area under the recall curve for transcription factors that rank less than 10 (Figure 3B) did not appear drastically different when ATAC-seq regions were required to overlap with EP300 or H3K27ac. Performance slightly decreased when using H3K4me1 or H3K4me3 data. The decrease in performance using H3K4me3 data is unsurprising given its known role in marking active promoters which may exclude enhancer regions that contain important transcription factor binding sites (Wamstad et al. 2014). We also examined the overlap of ATAC-seq with the three epigenomic signals marking active enhancers, EP300, H3K27ac, and H3K4me3 and found when selecting input sequences based on these criteria, the performance was similar to ATAC-seq and EP300 or ATAC-seq alone (Figure 3A-B). Using best performing choice of genomic-matched, shuffled, or enhancer background sequences, number of regions, and choice of stem cell or fibroblast to eliminate regions that were not cell type-specific, we found that H3K27ac, H3K4me1, or EP300 information did not significantly change the identification of reprogramming factors which is consistent with previous work suggesting that accessibility and H3K27ac have similar levels of accuracy in predicting enhancers validated by transgenic mouse assays (Fu et al. 2018). However, similar to the trend over all iterations of region selection the use of H3K4me3 resulted in marked decrease in recovery of transcription factors (Figure 3C). We also examined the particular transcription factors found with each histone mark and found no bias in transcription factors recovered by a particular epigenomic signal that was consistent across methods (Figure 3D).

**Figure 3.**
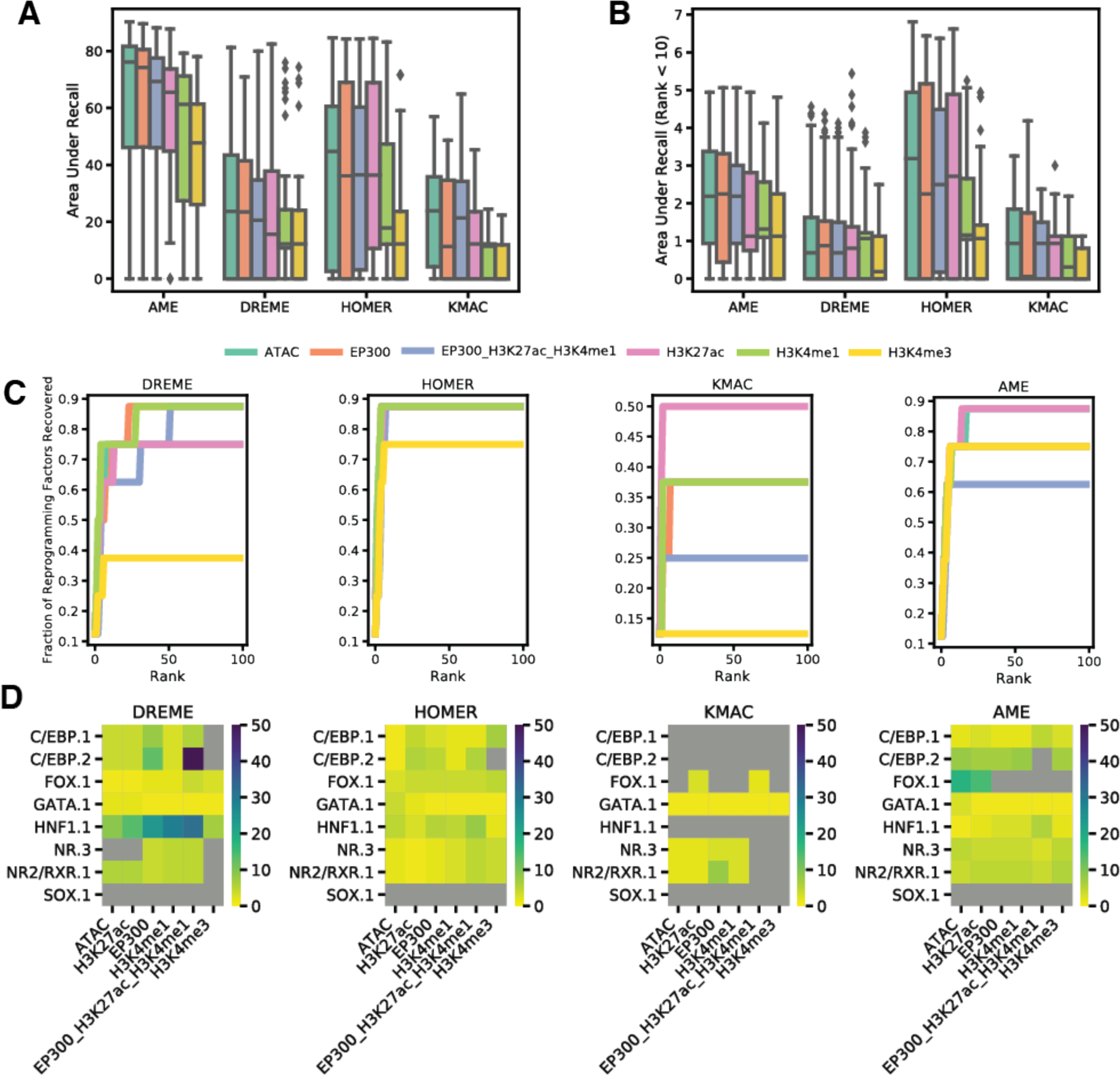
Use of histone mark and EP300 annotation does not significantly impact transcription factor recovery. A) Area under the recall curve for 4 methods (KMAC, AME, HOMER, and DREME) on hepatocyte transcription factor recovery shows an overall robust performance for all histone marks, except a slight dip in performance for H3K4me3 and overlap of EP300, H3K27ac, and H3K4me1. B) Area under the recall curve at rank less than 10 for 4 methods (KMAC, AME, HOMER, and DREME) on hepatocyte transcription factor recovery. C) Recall curves for the method with best area under the recall curve for rank less than 10 shows similar performance for ATAC, EP300, H3K27ac with lowered performance for H3K4me1, H3K4me3, and the overlap of all enhancer histone marks. D) Rank of each reprogramming transcription factor for each method and each genomic signal shows similar transcription factors recovered with similar rankings for all methods.

### Chromatin accessibility methods are highly effective for transcription factor recovery and significance ranking

We then evaluated all RNA-seq and ATAC-seq methods on their ability to identify known reprogramming factors. Overall, we find that chromatin accessibility methods outperform gene expression-based methods in terms of factor recall (Figure 4A). In particular, HOMER, AME, DeepAccess, and diffTF all robustly identify known reprogramming factors within the top 10 motifs. These trends hold across our 8 target cell types (Figure 4B). Looking at the performance as area under the recall curves for each cell type showed that AME, diffTF, and DeepAccess are more robustly able to identify a large number of reprogramming factors, likely because these methods place a prior on the discovery of motifs by using a transcription factor database (Figure 4C). When the recall curve is restricted to the top 10 ranked motifs, it shows that HOMER has impressive performance as a *de novo* motif discovery method in identifying reprogramming factors within its top enriched motifs (Figure 4D; Table S3). For 5 out of the 9 algorithms tested, using stem cells as the starting cell state resulted in a better recall of transcription factors over fibroblast though these performance differences were overall subtle (Figure S3; Fraction recovered at rank less than 10 – DREME: 0.42 (fibroblast), 0.44 (stem cell); AME: 0.55 (fibroblast), 0.68 (stem cell); KMAC: 0.21 (fibroblast), 0.24 (stem cell); HOMER 0.45 (fibroblast), 0.32 (stem cell); DeepAccess: 0.38 (fibroblast), 0.32 (stem cell); diffTF: 0.59 (fibroblast), 0.56 (stem cell); EBSeq 0.26 (fibroblast), 0.35 (stem cell); GarNet: 0.17 (fibroblast), 0.2 (stem cell)).

**Figure 4.**
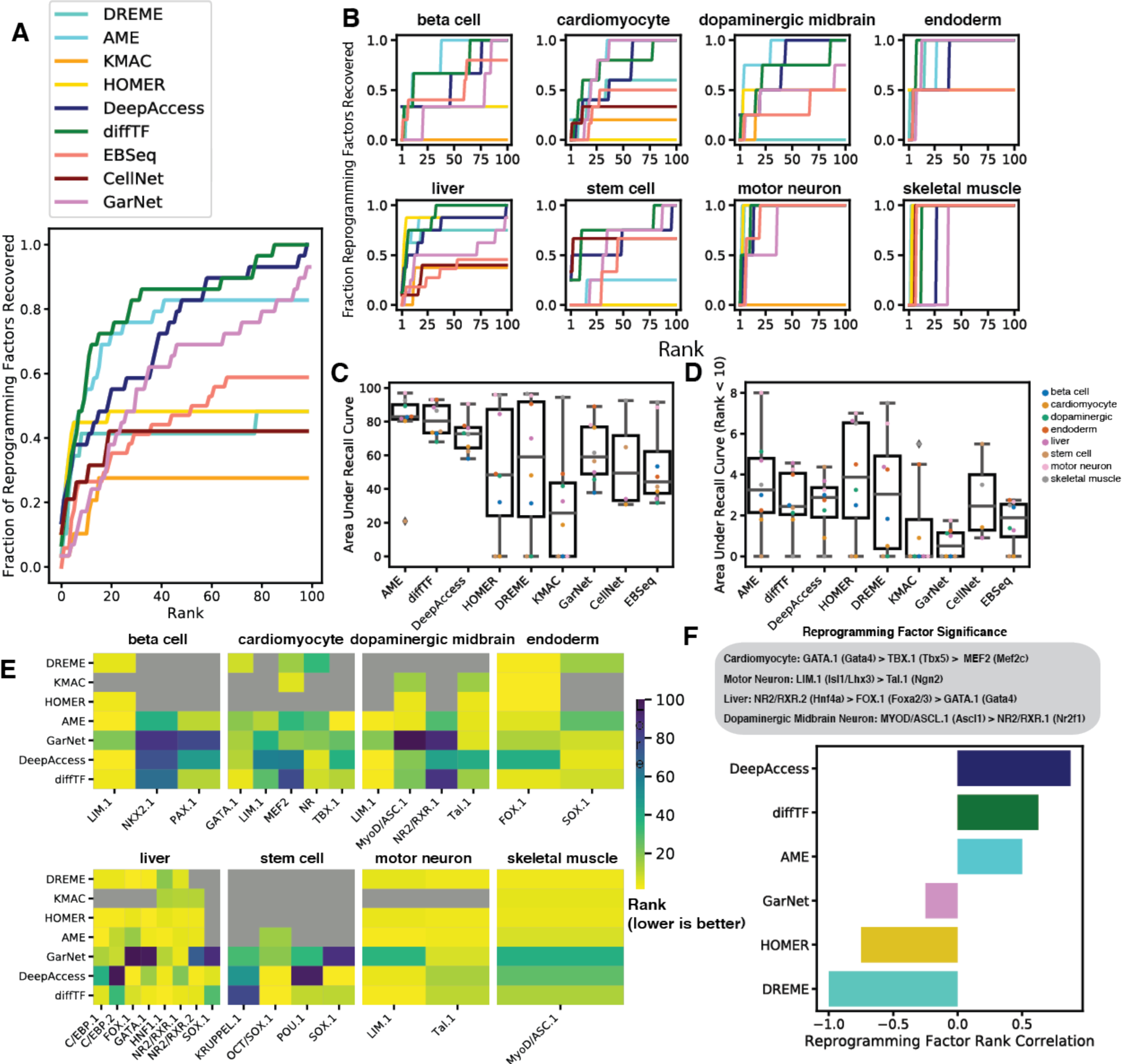
Complex chromatin methods are top performers for transcription factor recovery and significance ranking. A) Evaluation of 9 methods (2 entirely RNA-seq-based, 1 RNA-seq and ATAC-seq, 4 traditional ATAC-seq, 2 complex ATAC-seq) shows superior performance of AME, DeepAccess, and diffTF in transcription factor recovery. B) Evaluation of methods for each cell type shows superior methods generally perform well across all cell types. C) Area under recall curve for each method in each cell type shows AME, diffTF, and DeepAccess are able to robustly cover transcription factors. D) Area under recall curve at rank less than 10 for top performer for each method in each cell type shows HOMER is able to highly rank important transcription factors. E) Rank of reprogramming transcription factors for each method and each cell type. F) Transcription factors ordered by significance based on literature shows DeepAccess, diffTF, and AME are more likely to correctly rank the significance of transcription factor candidates.

Against our expectations, GarNet’s incorporation of gene expression with chromatin accessibility appeared to degrade rather than improve transcription factor recovery. There are several possible explanations for this observation that are not mutually exclusive: 1) by conditioning on proximity to genes, we eliminate distal accessible regions which have important regulatory function in reprogramming or accessible regions that may be cell type-specific but regulate genes that are not differential between starting and target cells, 2) transcription factor binding sites cannot be assumed to regulate their closest genes, or 3) transcription factor motif match score is not a good quantifier of impact of the transcription factor on gene expression. In order to test whether conditioning on proximity to genes had an impact on GarNet’s performance, we tested GarNet’s ability to identify transcription factors using 2kb, 10kb, or 100kb distance thresholds as limits to transcription factor:gene relationships. We found that the choice of distance threshold had little impact on GarNet’s performance (Figure S2).

When examining transcription factor rank within each cell type, it appeared that certain motifs were more robustly found across many methods, though only a few motifs were found by the best performer for all methods (Figure 4E). diffTF and DeepAccess were the only two methods that highly ranked the Oct4/Sox2 heterodimer motif, which is notable as a well-known transcription factor pair in both reprogramming (Yamamizu et al. 2013; Takahashi and Yamanaka 2006) and in manipulating nucleosomes to increase chromatin accessibility (Soufi et al. 2015). The disagreement in ranking between different methods led us to question whether methods were able to predict the relative significance of known reprogramming factors. From the literature, we found evidence of reprogramming factor significance for liver, pan-neuronal, and motor neuron by impact of over-expression of factors on reprogramming efficiency (Yamamizu et al. 2013; Simeonov and Uppal 2014) and evidence of significance in cardiomyocyte from tuned expression vector of Gata4, Tbx5, and Mef2c (Bai et al. 2015) as well as evidence of the significance of Gata4 in cardiomyocyte reprogramming from leave-one-out experiments (Jin et al. 2018). We then computed the correlation between true rank and observed rank for each method. Overall, we found that DeepAccess had the strongest correlation between true and observed rank (Figure 4F), with diffTF and AME ranking second and third.

## Discussion

Efficient and accurate transcription factor candidate prioritization for cellular reprogramming is an important unsolved problem. We evaluated eight complementary methods for this task on their ability to rediscover existing reprogramming protocols. These methods fell into three broad categories: gene expression-based, traditional epigenomic-based, and complex epigenomic-based.

Among the expression-based methods, CellNet represents a class of methods that use perturbation experiments to build a cell type-specific gene regulatory network. From these networks, hub genes are identified as potential targetable candidates for reprogramming. However, we found that while CellNet outperformed EBSeq for methods that use differential gene expression (Figure 4A), it was impossible to apply CellNet to new gene expression data with only few unperturbed replicates of the experiment. Another method that builds a regulatory network is GarNet, which links genes to proximal transcription factor binding sites within accessible genomic regions to build a transcription factor:gene regulatory network. GarNet then uses sparse linear regression to rank transcription factor candidates. While GarNet is a compelling approach as it combines gene expression information and accessibility information, we found that it did not outperform other chromatin accessibility methods (Figure 4A) which suggests that GarNet’s assumptions may not be optimal. For example, GarNet assumes that transcription factors within enhancers regulate their closest genes, and that factor binding strength is represented by PWM match score. Future research could apply GarNet using known enhancer:gene interactions from 3D interactome data and transcription factor binding strength from ChIP-seq to determine whether modifications to these assumptions can improve its performance.

We then compared all nine methods on their ability to recover transcription factors from eight cell types with successful differentiation protocols (Figure 1A). Overall, we found that AME, DeepAccess, and diffTF were able to outperform expression-based methods (Figure 4A-C). We found that AME was the top performer in reprogramming factor recovery when looking at all ranked motifs for each method (Figure 4A,C). However, when looking at only the top 10 ranked motifs, we found that HOMER was also very effective (Figure 4D). Methods such as AME, DeepAccess, GarNet, and diffTF rely on pre-existing motif databases that may be missing certain transcription factors entirely and cannot discover novel motifs such as a motif resulting from the binding of a heterodimer of two transcription factors. We also noticed a high variance for all methods in transcription factor recovery within the top 10 motifs (Figure 4D), which may be related to data quality but is difficult to disentangle from possible biological confounders like the developmental timepoint of the collected data or the complexity of the regulation of differentiation for different cell types.

Certain transcription factors were uniformly highly ranked across methods, while transcription factors such as Oct4/Sox2 motif were highly ranked only by the complex chromatin accessibility methods, diffTF and DeepAccess (Figure 4F). Observation of rank differences between methods lead us to ask whether certain methods were better at identifying the most significant transcription factor. We found evidence from the literature that supported ranking transcription factors within a reprogramming protocol, either through reprogramming efficiency of single transcription factor over-expression or through expression level optimization and leave-one-out experiments (Figure 4F). While metrics for reprogramming efficiency and specific cell type can make these comparisons difficult, significance ranking is an important additional metric as the transcription factors that are chosen to be included in a reprogramming protocol by ad-hoc methods may bias the results seen from evaluating recovery performance alone. diffTF and DeepAccess both were able to correctly rank the significance of transcription factors within reprogramming protocols (Figure 4F), indicating that there are advantages in using complex methods for identifying transcription factors from chromatin accessibility. Both methods also provide additional value beyond transcription factor discovery or enrichment. Predictive deep learning models like DeepAccess have been used to predict causal variants (Zhou and Troyanskaya 2015; Kelley et al. 2016, 2018), interpret complex genomic grammar (Koo et al. 2018; Avsec et al. 2020; Kim et al. 2020), and identify evolutionarily orthologous enhancers (Kelley 2020; Minnoye et al. 2020). In contrast with all other approaches, DeepAccess predictions for transcription factor effect come from designed genomic sequences containing transcription factor motifs. Therefore, DeepAccess can used to probe more complex regulatory grammar such as transcription factor combinations or spacing. Despite a tradeoff in computational expense of training neural networks, DeepAccess may be preferred if the user intends to investigate regulatory logic beyond transcription factor activity. In contrast, diffTF may desirable for its statistical approach if another goal is to be able to identify whether individual binding sites are differentially accessible.

While we found that expression-based methods were worse performers than accessibility-based methods for transcription factor recovery, we note that comparing performance between chromatin accessibility methods and gene expression methods is imperfect. We require perfect transcription factor matches when evaluating CellNet and EBSeq, while we require transcription factor family matches for GarNet, HOMER, AME, DREME, KMAC, diffTF, and DeepAccess. Ultimately, the best approach is likely to incorporate expression and accessibility data, but rather than taking GarNet’s approach to using expression to filter accessible regions and rank transcription factors, our results would suggest it is best to first use chromatin accessibility alone to select transcription factor families and then use gene expression to select differentially expressed transcription factors within each family.

In evaluating methods, the best methods only reached 50-60% recall of reprogramming factors within the top 10 motifs, indicating there is still room for methodological advancements. Single cell expression data may provide sufficient information for methods such as CellNet to be applied with less experimental burden. Methods like GarNet that use both expression and accessibility may require further exploration to determine optimal approaches to combine these data. Methods such as diffTF and GarNet that rely on input of transcription factor binding sites may suffer from issues in determining an appropriate binding cutoff based on PWM information alone, and could be combined with more complex binding prediction models like DeepBind (Alipanahi et al. 2015) to improve performance.

A limitation of our work is that we assumed there were no alternative factors to the ones present in the reference reprogramming protocols we utilized. This assumption ignores potentially better candidates that are experimentally uncharacterized. We examined whether there were any potentially novel reprogramming targets that were consistently ranked by HOMER, AME, DeepAccess, and diffTF (Table S4). Among these, we found NFI half-motif which was identified by 3 out of the 4 methods as a potential skeletal muscle regulator. Indeed, NFIX has been previously implicated in skeletal muscle development (Pistocchi et al. 2013; Messina et al. 2010). Hnf1 and Rfx were ranked by all methods within the top 10 motifs for beta pancreatic cells, with Hnf1 having a role in early progenitor development of endocrine cells (De Vas et al. 2015) and Rfx transcription factors playing an important role in adult pancreatic cell functional development (Ait-Lounis et al. 2010; Piccand et al. 2014). Experimental methods to evaluate transcription factors in parallel through transcription factor screens (Liu et al. 2018; Yang et al. 2019; Black et al. 2020; Genga et al. 2019; Ng et al. 2020; Nakatake et al. 2020) or high-throughput reporter assays for functional transcription factor activity such as chromatin accessibility (Hammelman et al. 2020) would allow us to expand our evaluation of methods to their ability to propose novel factors.

Overall, we found that chromatin accessibility-based methods recover the most known reprogramming transcription factors. We suggested methods for optimal region selection for traditional accessibility-based methods, and suggest that there are some performance benefits to using complex accessibility-based methods that perform more statistical or complex sequencing modeling of epigenomic data. We hope that our comprehensive evaluation of transcription factor reprogramming ranking and recovery will contribute to a basis for motif analysis procedures and a standard evaluation metric in the development of novel computational methods.

## Acknowledgements

We thank members of the Gifford lab and Wichterle lab for helpful discussions. We gratefully acknowledge funding from 1RO1HG008363 (D.K.G.), 1R01HG008754 (D.K.G.), 1R01NS109217 (D.K.G. & H.W.), R01NS116141 (H.W.), NINDS Postdoctoral NRSA Fellowship (F32NS105372) (T.P.), and National Science Foundation Graduate Research Fellowship (1122374) (J.H.).

## Contributions

Conceptualization, J.H., T.P, M.C., and D.K.G.; Methodology, J.H., T.P., M.C.; Software, J.H.; Formal Analysis, J.H.; Investigation, J.H.; Resources, D.K.G. & H.W.; Data Curation, J.H.; Writing - Original Draft, J.H.; Writing - Review & Editing, J.H., T.P., M.C., H.W., & D.K.G.; Visualization, J.H.; Supervision, D.K.G. & H.W.; Funding Acquisition, J.H., T.P., H.W. & D.K.G.

## Methods

### Data and Code Availability

The consensus mouse transcription factor motif database, shared mouse enhancer sequences, and a list of mouse transcription factors as well as a script for performing motif discovery with AME, DREME, HOMER, and KMAC is available at https://cgs.csail.mit.edu/ReprogrammingRecovery/.

Publicly available ATAC-seq and RNA-seq samples were downloaded as fastqs from Nucleotide Read Archive and processed as described in sections ATAC-seq processing and RNA-seq processing below. Uniformly processed gene count and peak files are also available at https://cgs.csail.mit.edu/ReprogrammingRecovery/.

### ATAC-seq processing

Reads were trimmed for adaptors and low-quality positions using Trimgalore (Cutadapt v0.6.2). Reads were aligned to the mouse genome (mm10) with bwa mem (v0.7.1.7) with default parameters. Properly paired mapped reads were filtered, and accessible regions were called using MACS2 (v2.2.7.1) with the parameters -f BAMPE -g mm -p 0.01 --shift -36 --extsize 73 -- nomodel --keep-dup all --call-summits. Accessible regions that overlapped genome blacklist regions (encode blacklist regions ENCSR535HHO) were excluded from downstream analysis.

### RNA-seq processing

Reads were trimmed for adaptors and low-quality positions using Trimgalore (Cutadapt v0.6.2). Reads were aligned to the mouse genome (mm10) and gene-level counts were quantified using RSEM (v1.3.0) rsem-calculate-expression using default parameters and STAR (v2.5.2b) for alignment.

### DeepAccess Training and Interpretation

We trained DeepAccess on data from ten cell types: stem cell, fibroblast, hepatocyte, endoderm, beta pancreatic cell, alpha pancreatic cell, cardiomyocyte, skeletal muscle, dopaminergic midbrain neuron, and spinal motor neuron using binary crossentropy loss (multitask classification) with 4,812,987 genomic regions for training: 3,133,509 regions were open in at least 1 cell type, and 1,679,478 regions were closed in all cell types (sampled with HOMER to match GC content % of accessible regions in genome). Chromosome 18 and chromosome 19 were held out for validation and training, respectively. To define training regions for DeepAccess, we generate 100bp genomic windows across the entire mouse genome. We define a region as accessible in a given cell type if more that 50% of the 100bp region overlaps a MACS2 accessible region from that cell type. We then compute the Differential Expected Pattern Effect (Hammelman and Gifford 2021) for all transcription factor motifs within the consensus database between the target and starting cell types.

### CellNet analysis

We ran CellNet using the provided mouse network and our fibroblast RNA-seq sample replicates as input to obtain a list of ranked genes.

### Transcription factor list curation and differential expression with EBSeq

After RSEM quantification, EBSeq was run with the RSEM wrapper rsem-run-ebseq. Genes were filtered to fdr p < 0.05. Analysis was limited to a list of 1,374 transcription factors derived from (Lambert et al. 2018), which were sorted by EBSeq estimates of change in expression between target and starting cell types. List of mouse transcription factors is available at https://cgs.csail.mit.edu/ReprogrammingRecovery/.

### GarNet analysis

PWMScan (v1.1.0) was used to identify transcription factor motif instances in the genome for the 107 consensus transcription factors. Since GarNet builds transcription factor:gene networks using proximity assignment, we built 3 GarNet networks where transcription factor influence was set to a threshold of 2kb, 10kb, or 100kb from transcription start sites. Then, GarNet was run for each starting cell type, target cell type pair with an input of differential expression (fold change) for genes with fdr < 0.05 for differential gene expression as estimated with EBSeq, and target cell type MACS2 accessible regions. Transcription factor rank was determined by the value of the slope.

### AME analysis

AME was run with default parameters with “--control --shuffle--” for a dinucleotide shuffled (default) background or enhancer background sequences were provided.

### DREME analysis

DREME was run with default parameters.

### HOMER analysis

For default background, HOMER was run using the command findMotifsGenome.pl target.bed mm10 target-homer -size given. For enhancer background, HOMER was run using the command findMotifs.pl target.fa fasta target-homer -fasta enhancer.fa.

### KMAC analysis

KMAC was run with the parameters --k_win 100 --k_min 4 --k_max 13 --t 1 --k_seqs 10000 -- k_top 10 --gap 4.

### diffTF analysis

PWMScan (v1.1.0) was used to identify transcription factor motif instances in the genome for the 107 consensus transcription factors. Then, diffTF was run with pairs of starting cell type, target cell type data sets with default parameters. Input was MACS2 accessible peaks and ATAC-seq aligned reads for sample replicates.

### *De novo* method transcription factor matching

To match transcription factor motifs to their best motif match for a known reprogramming transcription factor, we ran Tomtom (v5.0.5) with default parameters and set a threshold of a motif match with q-value < 0.05. The lowest rank (most enriched) motif that a given reprogramming transcription factor is assigned to that transcription factor.

### Software

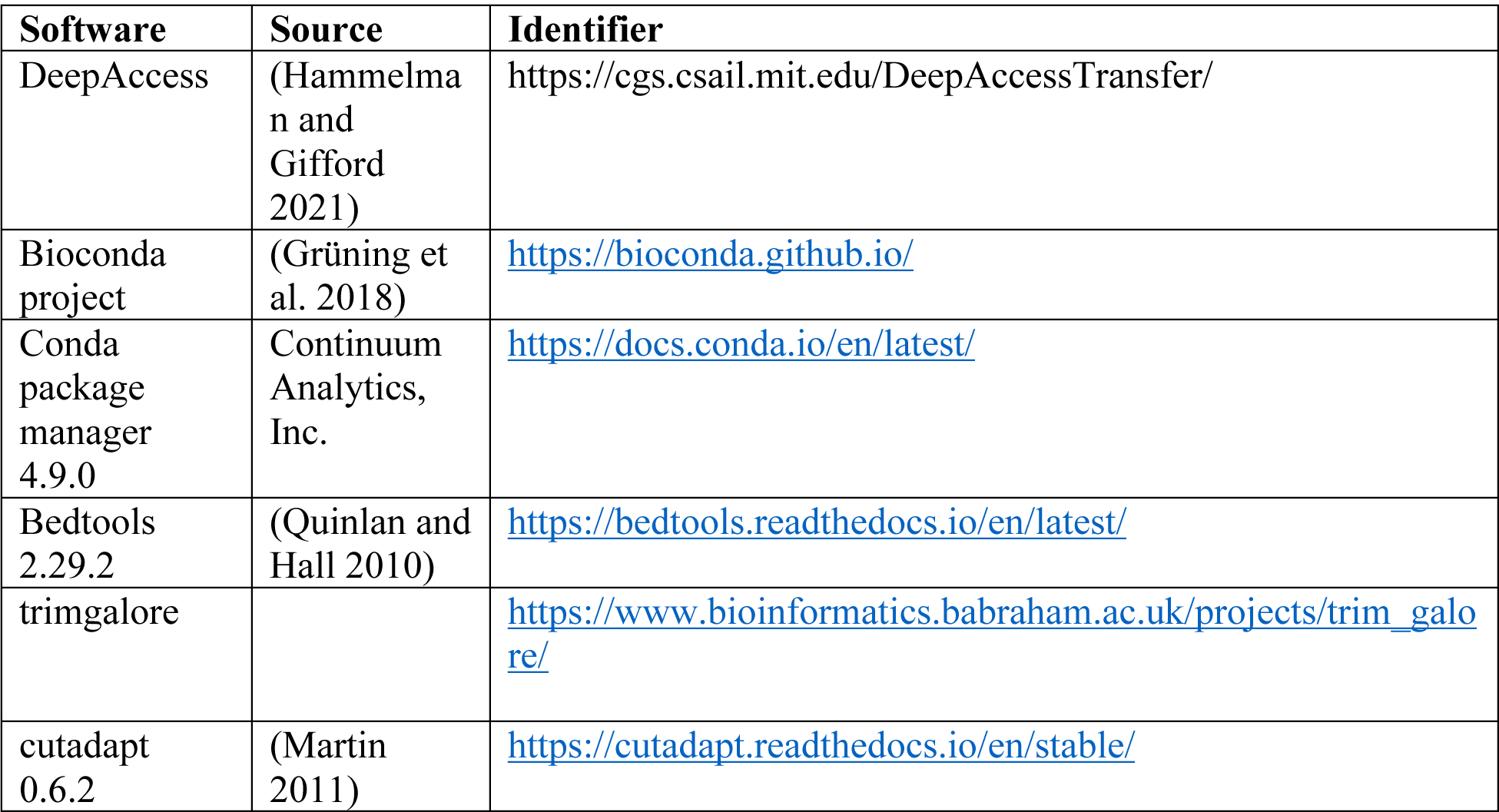
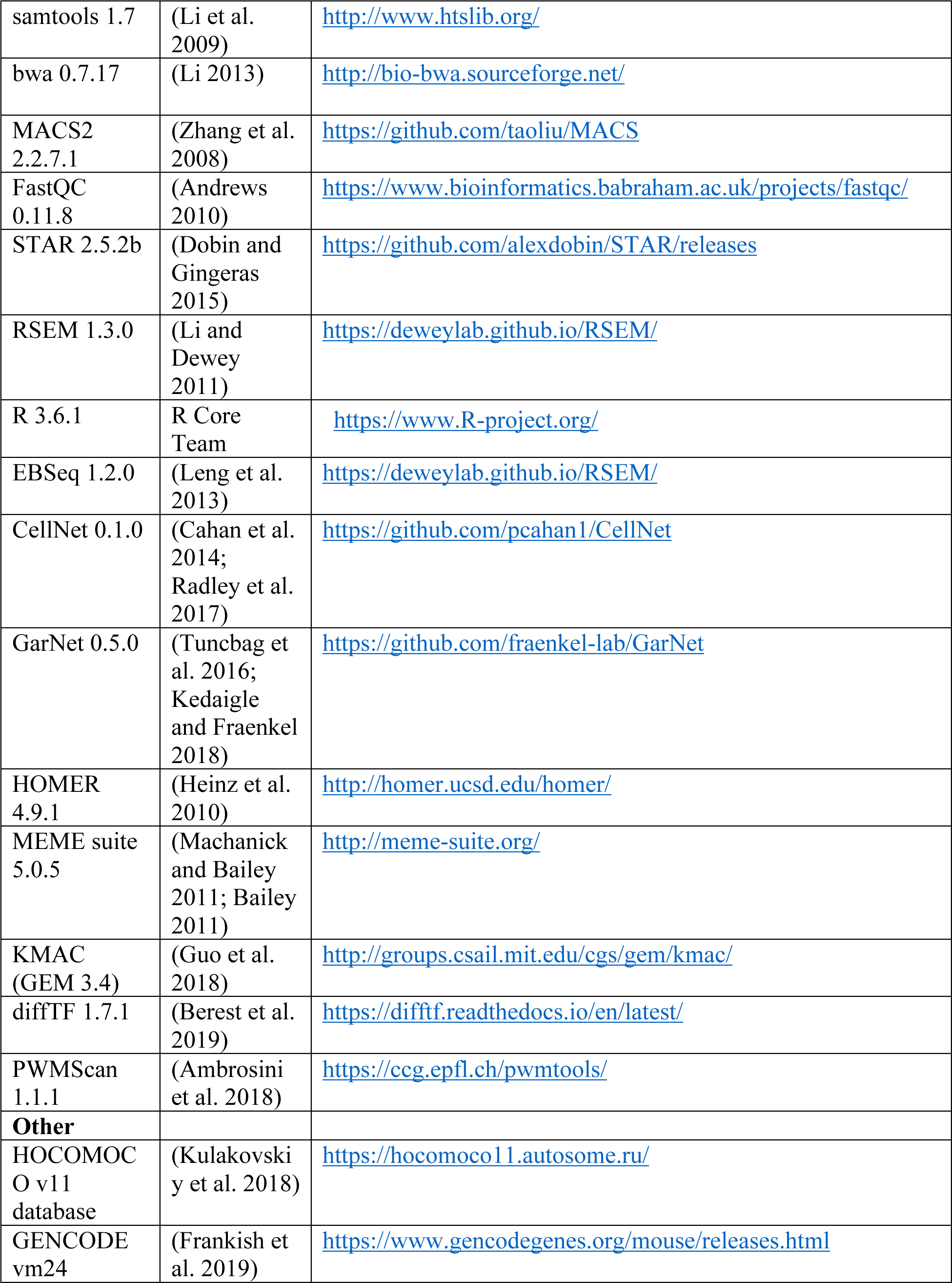

## References

Ait-Lounis A, Bonal C, Seguín-Estévez Q, Schmid CD, Bucher P, Herrera PL, Durand B, Meda P, Reith W. 2010. The transcription factor Rfx3 regulates beta-cell differentiation, function, and glucokinase expression. Diabetes 59: 1674–1685. https://pubmed.ncbi.nlm.nih.gov/20413507.

Alipanahi B, Delong A, Weirauch MT, Frey BJ. 2015. Predicting the sequence specificities of DNA- and RNA-binding proteins by deep learning. Nat Biotechnol 33: 831. http://dx.doi.org/10.1038/nbt.3300.

Ambrosini G, Groux R, Bucher P. 2018. PWMScan: a fast tool for scanning entire genomes with a position-specific weight matrix. Bioinformatics 34: 2483–2484.

Andrews S. 2010. FastQC: a quality control tool for high throughput sequence data.

Angermueller C, Clark SJ, Lee HJ, Macaulay IC, Teng MJ, Hu TX. 2016. Parallel single-cell sequencing links transcriptional and epigenetic heterogeneity. Nat Methods 13. http://dx.doi.org/10.1038/nmeth.3728.

Avsec Ž, Weilert M, Shrikumar A, Krueger S, Alexandari A, Dalal K, Fropf R, McAnany C, Gagneur J, Kundaje A, et al. 2020. Base-resolution models of transcription factor binding reveal soft motif syntax. bioRxiv 737981. http://biorxiv.org/content/early/2020/07/19/737981.abstract.

Bai F, Ho Lim C, Jia J, Santostefano K, Simmons C, Kasahara H, Wu W, Terada N, Jin S. 2015. Directed Differentiation of Embryonic Stem Cells Into Cardiomyocytes by Bacterial Injection of Defined Transcription Factors. Sci Rep 5: 15014. https://doi.org/10.1038/srep15014.

Bailey TL. 2011. DREME: motif discovery in transcription factor ChIP-seq data. Bioinformatics 27:1653–1659.

Berest I, Arnold C, Reyes-Palomares A, Palla G, Rasmussen KD, Giles H, Bruch P-M, Huber W, Dietrich S, Helin K. 2019. Quantification of differential transcription factor activity and multiomics-based classification into activators and repressors: diffTF. Cell Rep 29: 3147–3159.

Black JB, McCutcheon SR, Dube S, Barrera A, Klann TS, Rice GA, Adkar SS, Soderling SH, Reddy TE, Gersbach CA. 2020. Master Regulators and Cofactors of Human Neuronal Cell Fate Specification Identified by CRISPR Gene Activation Screens. Cell Rep 33. https://doi.org/10.1016/j.celrep.2020.108460.

Bonneau R, Reiss DJ, Shannon P, Facciotti M, Hood L, Baliga NS, Thorsson V. 2006. The Inferelator: an algorithm for learning parsimonious regulatory networks from systems-biology data sets de novo. Genome Biol 7: R36. https://doi.org/10.1186/gb-2006-7-5-r36.

Buenrostro J, Wu B, Chang H, Greenleaf W. 2015. ATAC-seq: A Method for Assaying Chromatin Accessibility Genome-Wide. Curr Protoc Mol Biol 109: 21.29.1–21.29.9. http://www.ncbi.nlm.nih.gov/pmc/articles/PMC4374986/.

Cahan P, Li H, Morris SA, Lummertz da Rocha E, Daley GQ, Collins JJ. 2014. CellNet: network biology applied to stem cell engineering. Cell 158: 903–915. https://pubmed.ncbi.nlm.nih.gov/25126793.

Cernilogar FM, Hasenoder S, Wang Z, Scheibner K, Burtscher I, Sterr M, Smialowski P, Groh S, Evenroed IM, Gilfillan GD, et al. 2019. Pre-marked chromatin and transcription factor cobinding shape the pioneering activity of Foxa2. Nucleic Acids Res 47: 9069–9086.

Corces MR, Trevino AE, Hamilton EG, Greenside PG, Sinnott-Armstrong NA, Vesuna S, Satpathy AT, Rubin AJ, Montine KS, Wu B. 2017. An improved ATAC-seq protocol reduces background and enables interrogation of frozen tissues. Nat Methods 14: 959–962.

D’Alessio AC, Fan ZP, Wert KJ, Baranov P, Cohen MA, Saini JS, Cohick E, Charniga C, Dadon D, Hannett NM, et al. 2015. A Systematic Approach to Identify Candidate Transcription Factors that Control Cell Identity. Stem Cell Reports 5: 763–775. http://www.sciencedirect.com/science/article/pii/S2213671115002787.

De Vas MG, Kopp JL, Heliot C, Sander M, Cereghini S, Haumaitre C. 2015. Hnf1b controls pancreas morphogenesis and the generation of Ngn3+ endocrine progenitors. Development 142: 871–882. https://pubmed.ncbi.nlm.nih.gov/25715395.

Dixit A, Parnas O, Li B, Chen J, Fulco CP, Jerby-Arnon L, Marjanovic ND, Dionne D, Burks T, Raychowdhury R, et al. 2016. Perturb-Seq: Dissecting Molecular Circuits with Scalable Single-Cell RNA Profiling of Pooled Genetic Screens. Cell 167: 1853–1866.e17. http://linkinghub.elsevier.com/retrieve/pii/S0092867416316105 (Accessed May 24, 2017).

Dobin A, Gingeras TR. 2015. Mapping RNA□seq reads with STAR. Curr Protoc Bioinforma 51: 11–14.

Frankish A, Diekhans M, Ferreira A-M, Johnson R, Jungreis I, Loveland J, Mudge JM, Sisu C, Wright J, Armstrong J. 2019. GENCODE reference annotation for the human and mouse genomes. Nucleic Acids Res 47: D766–D773.

Frey BJ, Dueck D. 2007. Clustering by passing messages between data points. Science (80-) 315: 972–976.

Fu S, Wang Q, Moore JE, Purcaro MJ, Pratt HE, Fan K, Gu C, Jiang C, Zhu R, Kundaje A, et al. 2018. Differential analysis of chromatin accessibility and histone modifications for predicting mouse developmental enhancers. Nucleic Acids Res 46: 11184–11201. https://doi.org/10.1093/nar/gky753.

Genga RMJ, Kernfeld EM, Parsi KM, Parsons TJ, Ziller MJ, Maehr R. 2019. Single-Cell RNA-Sequencing-Based CRISPRi Screening Resolves Molecular Drivers of Early Human Endoderm Development. Cell Rep 27: 708–718.e10. http://www.sciencedirect.com/science/article/pii/S2211124719304061.

Grün D, Lyubimova A, Kester L, Wiebrands K, Basak O, Sasaki N. 2015. Single-cell messenger RNA sequencing reveals rare intestinal cell types. Nature 525. http://dx.doi.org/10.1038/nature14966.

Grüning B, Chilton J, Köster J, Dale R, Soranzo N, van den Beek M, Goecks J, Backofen R, Nekrutenko A, Taylor J. 2018. Practical computational reproducibility in the life sciences. Cell Syst 6: 631–635.

Guo Y, Tian K, Zeng H, Guo X, Gifford DK. 2018. A novel k-mer set memory (KSM) motif representation improves regulatory variant prediction. Genome Res.

Gupta S, Stamatoyannopoulos JA, Bailey TL, Noble WS. 2007. Quantifying similarity between motifs. Genome Biol 8: R24. https://doi.org/10.1186/gb-2007-8-2-r24.

Habib N, Li Y, Heidenreich M, Swiech L, Avraham-Davidi I, Trombetta JJ, Hession C, Zhang F, Regev A. 2016. Div-Seq: Single-nucleus RNA-Seq reveals dynamics of rare adult newborn neurons. Science (80-) 353: 925–928.

Hammelman J, Gifford DK. 2021. Discovering differential genome sequence activity with interpretable and efficient deep learning. bioRxiv.

Hammelman J, Krismer K, Banerjee B, Gifford DK, Sherwood RI. 2020. Identification of determinants of differential chromatin accessibility through a massively parallel genome-integrated reporter assay. Genome Res 30.

Heinäniemi M, Nykter M, Kramer R, Wienecke-Baldacchino A, Sinkkonen L, Zhou JX, Kreisberg R, Kauffman SA, Huang S, Shmulevich I. 2013. Gene-pair expression signatures reveal lineage control. Nat Methods 10: 577–583.

Heinz S, Benner C, Spann N, Bertolino E, Lin YC, Laslo P, Cheng JX, Murre C, Singh H, Glass CK. 2010. Simple combinations of lineage-determining transcription factors prime cis-regulatory elements required for macrophage and B cell identities. Mol Cell 38: 576–589.

Ichida JK, Blanchard J, Lam K, Son EY, Chung JE, Egli D, Loh KM, Carter AC, Di Giorgio FP, Koszka K. 2009. A small-molecule inhibitor of Tgf-ß signaling replaces Sox2 in reprogramming by inducing Nanog. Cell Stem Cell 5: 491–503.

Jin Y, Liu Y, Li Z, Santostefano K, Shi J, Zhang X, Wu D, Cheng Z, Wu W, Terada N. 2018. Enhanced differentiation of human pluripotent stem cells into cardiomyocytes by bacteria-mediated transcription factors delivery. PLoS One 13: e0194895.

Kedaigle AJ, Fraenkel E. 2018. Discovering altered regulation and signaling through networkbased integration of transcriptomic, epigenomic, and proteomic tumor data. In Cancer Systems Biology, pp. 13–26, Springer.

Kelley DR. 2020. Cross-species regulatory sequence activity prediction. PLoS Comput Biol 16: e1008050.

Kelley DR, Reshef YA, Bileschi M, Belanger D, McLean CY, Snoek J. 2018. Sequential regulatory activity prediction across chromosomes with convolutional neural networks. Genome Res 28: 739–750.

Kelley DR, Snoek J, Rinn JL. 2016. Basset: Learning the regulatory code of the accessible genome with deep convolutional neural networks. Genome Res 26: 990–999.

Kim D, Risca V, Reynolds D, Chappell J, Rubin A, Jung N, Donohue L, Kathiria A, Shi M, Zhao Z, et al. 2020. The dynamic, combinatorial cis-regulatory lexicon of epidermal differentiation. bioRxiv 2020.10.16.342857. http://biorxiv.org/content/early/2020/10/18/2020.10.16.342857.abstract.

Koo PK, Anand P, Paul SB, Eddy SR. 2018. Inferring Sequence-Structure Preferences of RNA-Binding Proteins with Convolutional Residual Networks. bioRxiv 418459.

Kulakovskiy I V, Vorontsov IE, Yevshin IS, Sharipov RN, Fedorova AD, Rumynskiy EI, Medvedeva YA, Magana-Mora A, Bajic VB, Papatsenko DA, et al. 2018. HOCOMOCO: towards a complete collection of transcription factor binding models for human and mouse via large-scale ChIP-Seq analysis. Nucleic Acids Res 46: D252–D259. https://doi.org/10.1093/nar/gkx1106.

Lake BB, Chen S, Sos BC, Fan J, Kaeser GE, Yung YC, Duong TE, Gao D, Chun J, Kharchenko P V, et al. 2018. Integrative single-cell analysis of transcriptional and epigenetic states in the human adult brain. Nat Biotechnol 36: 70–80. https://pubmed.ncbi.nlm.nih.gov/29227469.

Lambert SA, Jolma A, Campitelli LF, Das PK, Yin Y, Albu M, Chen X, Taipale J, Hughes TR, Weirauch MT. 2018. The human transcription factors. Cell 172: 650–665.

Lang AH, Li H, Collins JJ, Mehta P. 2014. Epigenetic landscapes explain partially reprogrammed cells and identify key reprogramming genes. PLoS Comput Biol 10: e1003734.

Leng N, Dawson JA, Thomson JA, Ruotti V, Rissman AI, Smits BMG, Haag JD, Gould MN, Stewart RM, Kendziorski C. 2013. EBSeq: an empirical Bayes hierarchical model for inference in RNA-seq experiments. Bioinformatics 29: 1035–1043. https://doi.org/10.1093/bioinformatics/btt087.

Li B, Dewey CN. 2011. RSEM: accurate transcript quantification from RNA-Seq data with or without a reference genome. BMC Bioinformatics 12. http://dx.doi.org/10.1186/1471-2105-12-323.

Li H. 2013. Aligning sequence reads, clone sequences and assembly contigs with BWA-MEM. arXiv Prepr arXiv13033997.

Li H, Handsaker B, Wysoker A, Fennell T, Ruan J, Homer N, Marth G, Abecasis G, Durbin R. 2009. The sequence alignment/map format and SAMtools. Bioinformatics 25: 2078–2079.

Liu C, Wang M, Wei X, Wu L, Xu J, Dai X, Xia J, Cheng M, Yuan Y, Zhang P, et al. 2019. An ATAC-seq atlas of chromatin accessibility in mouse tissues. Sci Data 6: 65. https://doi.org/10.1038/s41597-019-0071-0.

Liu Y, Yu C, Daley TP, Wang F, Cao WS, Bhate S, Lin X, Still II C, Liu H, Zhao D. 2018. CRISPR activation screens systematically identify factors that drive neuronal fate and reprogramming. Cell Stem Cell 23: 758–771.

Machanick P, Bailey TL. 2011. MEME-ChIP: motif analysis of large DNA datasets. Bioinformatics 27: 1696–1697.

Marson A, Foreman R, Chevalier B, Bilodeau S, Kahn M, Young RA, Jaenisch R. 2008. Wnt signaling promotes reprogramming of somatic cells to pluripotency. Cell Stem Cell 3: 132.

Martin M. 2011. Cutadapt removes adapter sequences from high-throughput sequencing reads. EMBnet J 17: 10–12.

Mazzoni EO, Mahony S, Closser M, Morrison CA, Nedelec S, Williams DJ. 2013. Synergistic binding of transcription factors to cell-specific enhancers programs motor neuron identity. Nat Neurosci 16. http://dx.doi.org/10.1038/nn.3467.

Messina G, Biressi S, Monteverde S, Magli A, Cassano M, Perani L, Roncaglia E, Tagliafico E, Starnes L, Campbell CE. 2010. Nfix regulates fetal-specific transcription in developing skeletal muscle. Cell 140: 554–566.

Minnoye L, Taskiran II, Mauduit D, Fazio M, Van Aerschot L, Hulselmans G, Christiaens V, Makhzami S, Seltenhammer M, Karras P. 2020. Cross-species analysis of enhancer logic using deep learning. Genome Res gr-260844.

Miraldi ER, Pokrovskii M, Watters A, Castro DM, De Veaux N, Hall JA, Lee J-Y, Ciofani M, Madar A, Carriero N. 2019. Leveraging chromatin accessibility for transcriptional regulatory network inference in T Helper 17 Cells. Genome Res 29: 449–463.

Morris SA, Cahan P, Li H, Zhao AM, San Roman AK, Shivdasani RA, Collins JJ, Daley GQ. 2014. Dissecting engineered cell types and enhancing cell fate conversion via CellNet. Cell 158:889–902.

Nakatake Y, Ko SBH, Sharov AA, Wakabayashi S, Murakami M, Sakota M, Chikazawa N, Ookura C, Sato S, Ito N, et al. 2020. Generation and Profiling of 2,135 Human ESC Lines for the Systematic Analyses of Cell States Perturbed by Inducing Single Transcription Factors. Cell Rep 31: 107655. http://www.sciencedirect.com/science/article/pii/S2211124720306082.

Ng AHM, Khoshakhlagh P, Rojo Arias JE, Pasquini G, Wang K, Swiersy A, Shipman SL, Appleton E, Kiaee K, Kohman RE, et al. 2020. A comprehensive library of human transcription factors for cell fate engineering. Nat Biotechnol. https://doi.org/10.1038/s41587-020-0742-6.

Oh Y, Jang J. 2019. Directed Differentiation of Pluripotent Stem Cells by Trascription Factors. Mol Cells.

Pellegrino M, Sciambi A, Yates JL, Mast JD, Silver C, Eastburn DJ. 2016. RNA-Seq following PCR-based sorting reveals rare cell transcriptional signatures. BMC Genomics 17: 1–12.

Piccand J, Strasser P, Hodson DJ, Meunier A, Ye T, Keime C, Birling M-C, Rutter GA, Gradwohl G. 2014. Rfx6 maintains the functional identity of adult pancreatic ß cells. Cell Rep 9: 2219–2232. https://pubmed.ncbi.nlm.nih.gov/25497096.

Pijuan-Sala B, Griffiths JA, Guibentif C, Hiscock TW, Jawaid W, Calero-Nieto FJ, Mulas C, Ibarra-Soria X, Tyser RC V, Ho DLL, et al. 2019. A single-cell molecular map of mouse gastrulation and early organogenesis. Nature 566: 490–495. https://doi.org/10.1038/s41586-019-0933-9.

Pijuan-Sala B, Wilson NK, Xia J, Hou X, Hannah RL, Kinston S, Calero-Nieto FJ, Poirion O, Preissl S, Liu F, et al. 2020. Single-cell chromatin accessibility maps reveal regulatory programs driving early mouse organogenesis. Nat Cell Biol 22: 487–497. https://doi.org/10.1038/s41556-020-0489-9.

Pistocchi A, Gaudenzi G, Foglia E, Monteverde S, Moreno-Fortuny A, Pianca A, Cossu G, Cotelli F, Messina G. 2013. Conserved and divergent functions of Nfix in skeletal muscle development during vertebrate evolution. Development 140: 1528–1536.

Quinlan AR, Hall IM. 2010. BEDTools: a flexible suite of utilities for comparing genomic features. Bioinformatics 26: 841–842.

Rackham OJL, Firas J, Fang H, Oates ME, Holmes ML, Knaupp AS, Suzuki H, Nefzger CM, Daub CO, Shin JW. 2016. A predictive computational framework for direct reprogramming between human cell types. Nat Genet 48: 331.

Radley AH, Schwab RM, Tan Y, Kim J, Lo EKW, Cahan P. 2017. Assessment of engineered cells using CellNet and RNA-seq. Nat Protoc 12: 1089–1102. https://doi.org/10.1038/nprot.2017.022.

Rai V, Quang DX, Erdos MR, Cusanovich DA, Daza RM, Narisu N, Zou LS, Didion JP, Guan Y, Shendure J. 2020. Single-cell ATAC-Seq in human pancreatic islets and deep learning upscaling of rare cells reveals cell-specific type 2 diabetes regulatory signatures. Mol Metab 32:109–121.

Roost MS, Van Iperen L, Ariyurek Y, Buermans HP, Arindrarto W, Devalla HD, Passier R, Mummery CL, Carlotti F, De Koning EJP. 2015. KeyGenes, a tool to probe tissue differentiation using a human fetal transcriptional atlas. Stem cell reports 4: 1112–1124.

Sasagawa Y, Nikaido I, Hayashi T, Danno H, Uno KD, Imai T, Ueda HR. 2013. Quartz-Seq: a highly reproducible and sensitive single-cell RNA-Seq reveals non-genetic gene expression heterogeneity. Genome Biol 14. http://dx.doi.org/10.1186/gb-2013-14-4-r31.

Satpathy AT, Granja JM, Yost KE, Qi Y, Meschi F, McDermott GP, Olsen BN, Mumbach MR, Pierce SE, Corces MR, et al. 2019. Massively parallel single-cell chromatin landscapes of human immune cell development and intratumoral T cell exhaustion. Nat Biotechnol 37: 925–936. https://doi.org/10.1038/s41587-019-0206-z.

Sharma N, Flaherty K, Lezgiyeva K, Wagner DE, Klein AM, Ginty DD. 2020. The emergence of transcriptional identity in somatosensory neurons. Nature 577: 392–398. https://doi.org/10.1038/s41586-019-1900-1.

Shen Y, Yue F, McCleary DF, Ye Z, Edsall L, Kuan S, Wagner U, Dixon J, Lee L, Lobanenkov V V. 2012. A map of the cis-regulatory sequences in the mouse genome. Nature 488: 116–120.

Simeonov KP, Uppal H. 2014. Direct Reprogramming of Human Fibroblasts to Hepatocyte-Like Cells by Synthetic Modified mRNAs. PLoS One 9: e100134. https://doi.org/10.1371/journal.pone.0100134.

Soufi A, Garcia MF, Jaroszewicz A, Osman N, Pellegrini M, Zaret KS. 2015. Pioneer transcription factors target partial DNA motifs on nucleosomes to initiate reprogramming. Cell 161: 555–568. https://www.ncbi.nlm.nih.gov/pubmed/25892221.

Takahashi K, Yamanaka S. 2006. Induction of Pluripotent Stem Cells from Mouse Embryonic and Adult Fibroblast Cultures by Defined Factors. Cell 126: 663–676. http://www.sciencedirect.com/science/article/pii/S0092867406009767.

Tuncbag N, Gosline SJC, Kedaigle A, Soltis AR, Gitter A, Fraenkel E. 2016. Network-based interpretation of diverse high-throughput datasets through the omics integrator software package. PLoS Comput Biol 12: e1004879.

Wamstad JA, Wang X, Demuren OO, Boyer LA. 2014. Distal enhancers: new insights into heart development and disease. Trends Cell Biol 24: 294–302. https://www.sciencedirect.com/science/article/pii/S096289241300189X.

Whitington T, Frith MC, Johnson J, Bailey TL. 2011. Inferring transcription factor complexes from ChIP-seq data. Nucleic Acids Res 39: e98–e98. https://doi.org/10.1093/nar/gkr341.

Wichterle H, Lieberam I, Porter JA, Jessell TM. 2002. Directed differentiation of embryonic stem cells into motor neurons. Cell 110: 385–397.

Yamamizu K, Piao Y, Sharov AA, Zsiros V, Yu H, Nakazawa K, Schlessinger D, Ko MSH. 2013. Identification of transcription factors for lineage-specific ESC differentiation. Stem cell reports 1: 545–559.

Yang J, Rajan SS, Friedrich MJ, Lan G, Zou X, Ponstingl H, Garyfallos DA, Liu P, Bradley A, Metzakopian E. 2019. Genome-scale CRISPRa screen identifies novel factors for cellular reprogramming. Stem cell reports 12: 757–771.

Zhang Y, Liu T, Meyer CA, Eeckhoute J, Johnson DS, Bernstein BE, Nusbaum C, Myers RM, Brown M, Li W, et al. 2008. Model-based analysis of ChIP-Seq (MACS). Genome Biol 9: R137.

Zhou J, Troyanskaya OG. 2015. Predicting effects of noncoding variants with deep learningbased sequence model. Nat Methods 12: 931–934. http://dx.doi.org/10.1038/nmeth.3547.

