## Supplemental Figures and Legends for "Ranking Reprogramming Factors for Directed Differentiation"

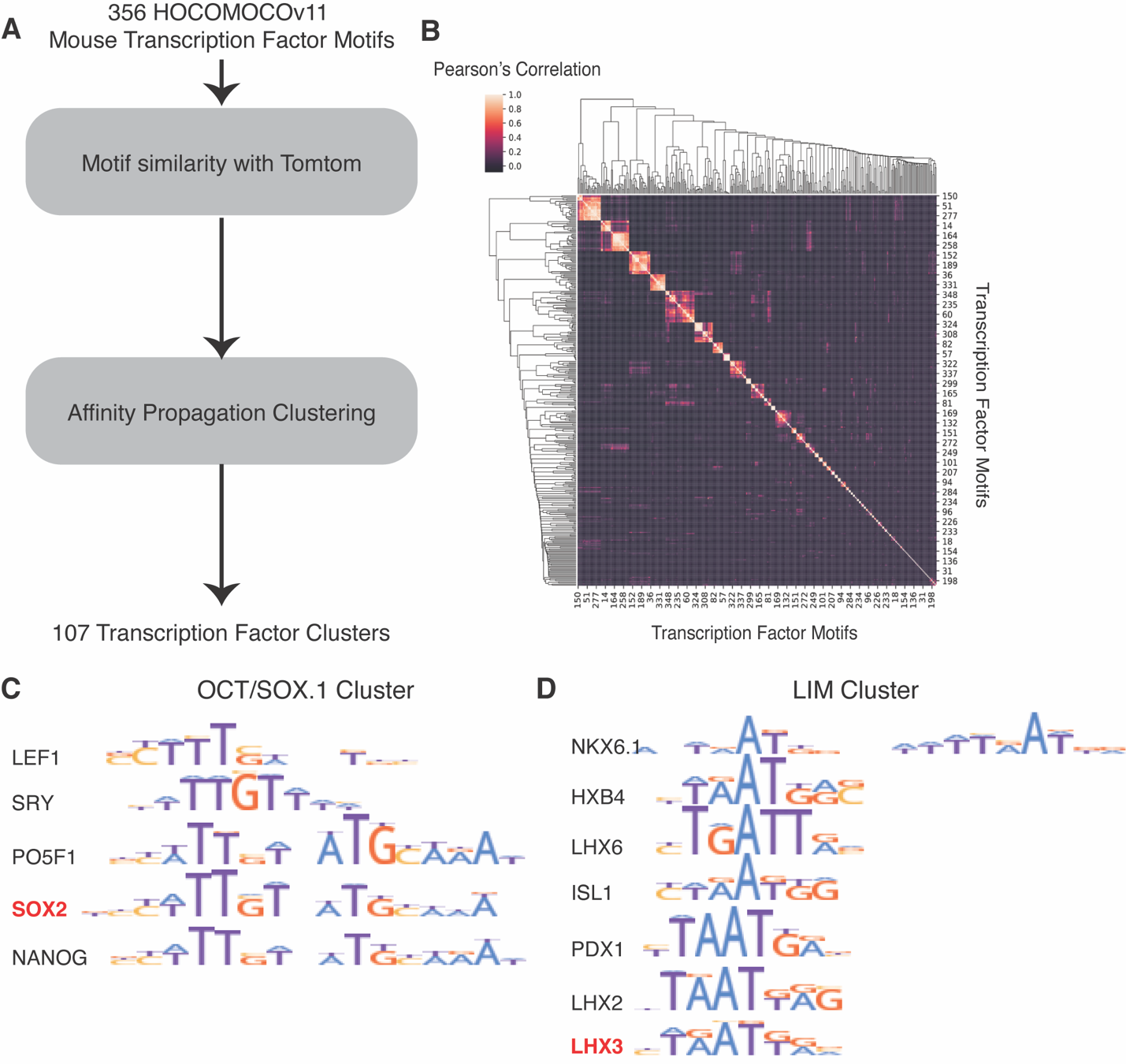

Supplemental Figure 1. A consensus database of 107 transcription factor motifs. A) HOCOMOCO v11 mouse transcription factor core motif database is used as input. Motif PWM similarity to the HOCOMOCO database is computed using Tomtom. B) For each pair of motifs, Pearson correlation between Tomtom scores is computed, resulting in a symmetric correlation matrix. Affinity propagation clustering is applied to the correlation matrix, resulting in 107 clusters of transcription factor motifs with one motif being selected as the representative motif of the cluster. C) Cluster representing OCT/SOX heterodimer-like motifs with SOX2 motif selected as the representative. D) Cluster representing LIM-like motifs with LHX3 motif selected as the representative.

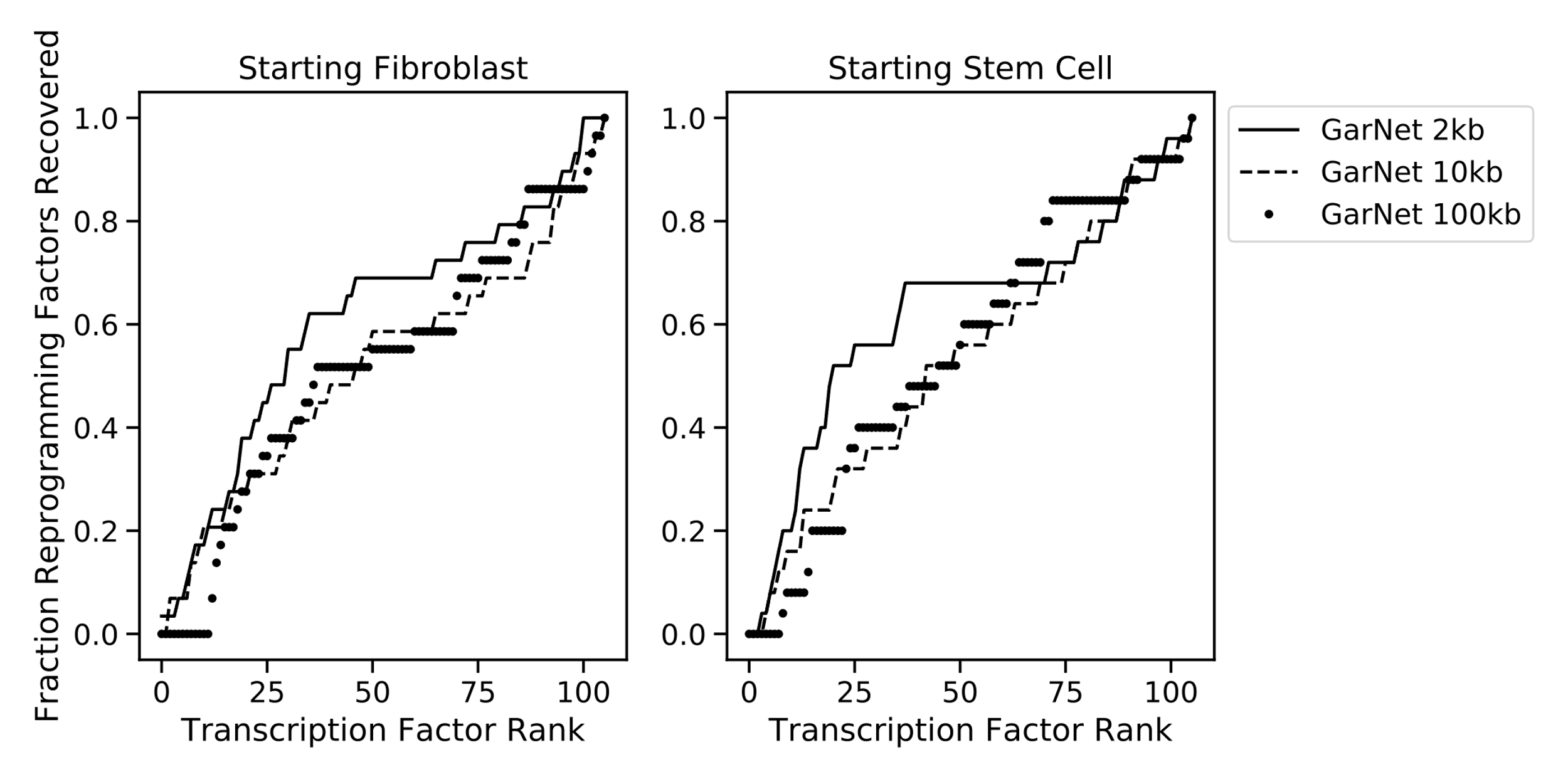

Supplemental Figure 2. GarNet fraction of reprogramming factors recovered with 2kb, 10kb, and 100kb thresholds for maximum distance between transcription factor binding site and gene transcription factor start site.

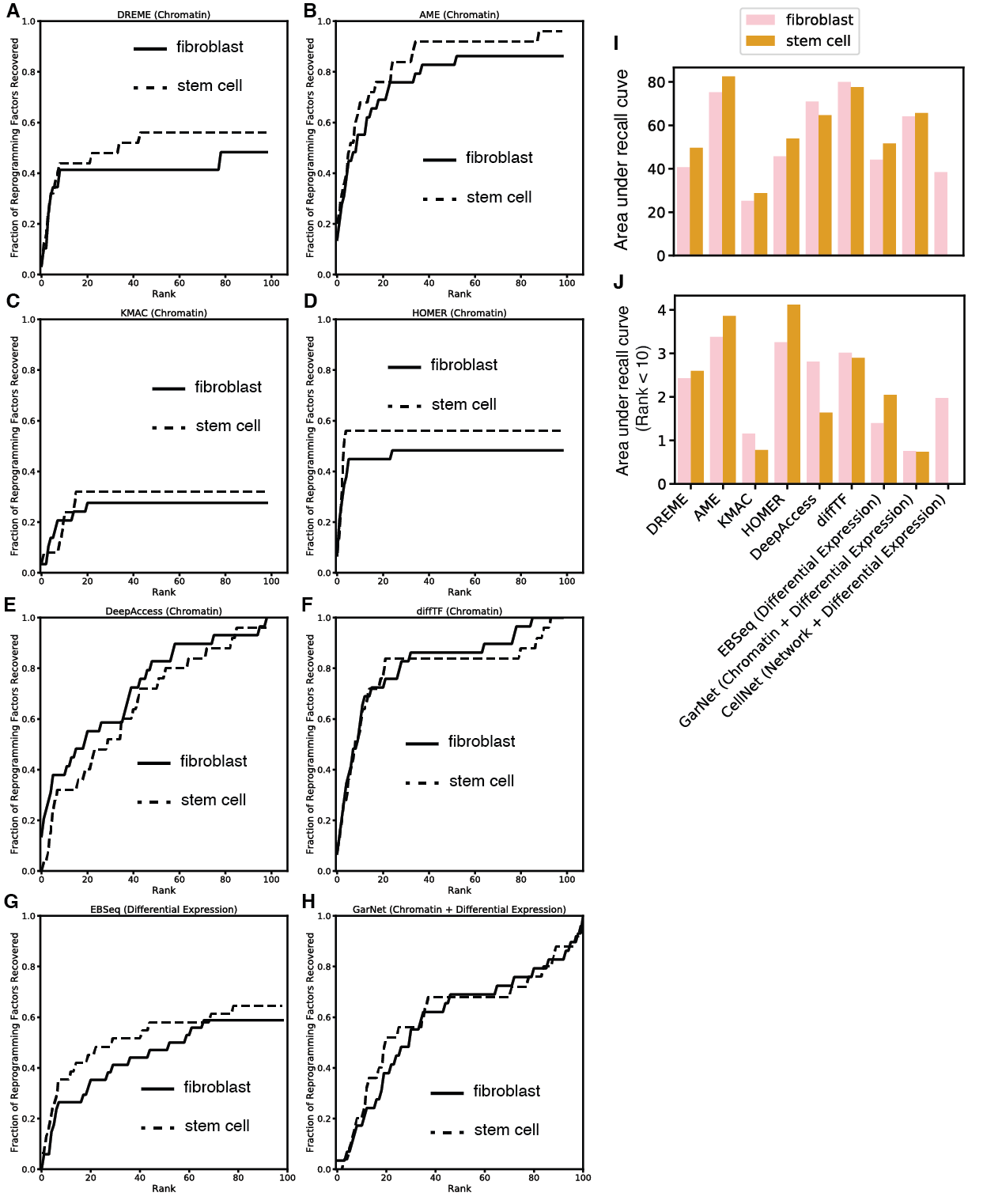

Supplemental Figure 3. Methods overall perform equally well using fibroblast or stem cell as starting cell state. A-H) Rank recall curves over all cell types for each algorithm using fibroblast or stem cell background. I) Area under recall curve for each method for fibroblast or stem cell source cells and J) area under recall curve at rank less than 10 for each method for fibroblast or stem cell source cells.

Table S1. Samples and accession numbers for RNA-seq and ATAC-seq data used for reprogramming factor recovery. All samples with the exception of endoderm were derived from primary cells or tissue.

Table S2. Area under the rank-recall curves by cell type for all chromatin accessibility-based methods.

Table S3. Area under the rank-recall curves for rank less than 10 for RNA and chromatin accessibility-based methods.

Table S4. Top 10 motifs ranked for reprogramming each cell type for methods diffTF, HOMER, AME, and DeepAccess.
